# All detectable ancient whole-genome duplications involve hybridization

**DOI:** 10.64898/2026.08.25.747000

**Authors:** Michelle L. Gaynor, Keyi Feng, Douglas E. Soltis, Pamela S. Soltis, Stephen A. Smith

**Affiliations:** Department of Ecology and Evolutionary Biology, University of Michigan, Ann Arbor, 48109 MI, USA; Florida Museum of Natural History, University of Florida, Gainesville, FL, 32611 USA; Department of Biology, University of Central Florida, Orlando, FL, 32816 USA; Department of Biology, University of Florida, Gainesville, FL, 32611 USA

**Keywords:** Polyploidy, Paleopolyploidy, Gene-divergence

## Abstract

Whole-genome duplication (WGD), or polyploidy, is a major evolutionary force throughout the tree of life. Ancient WGDs are commonly inferred from the distribution of synonymous substitutions per synonymous site (*K_s_*) between duplicated genes. Here, we argue that the standard interpretation of these distributions is incomplete. Using haplotype-phased genome assemblies spanning canonical autopolyploid and allopolyploid systems, we find that *K_s_* accumulates primarily between gene copies on non-recombining chromosomes. Thus, *K_s_*-based WGD detection depends not simply on WGD, but on whether duplicated copies evolve independently. This distinction reframes many inferences of ancient WGD as signatures of hybridization and the evolution of meiotic isolation and alters how we interpret the origin, detectability, and evolutionary consequences of polyploidy across the tree of life.

**Significance:** Whole-genome duplication (WGD) is prevalent across the tree of life, and its signatures appear in nearly every sequenced genome. Here we show that the predominant gene-based approach for detecting ancient WGD events is biased towards allopolyploid events arising through hybridization. The current approach is poorly suited to detecting WGD that occurs within a single species, also known as autopolyploidy. These findings suggest we likely underestimate the frequency of ancient polyploids. Furthermore, most signals of ancient WGD may therefore record not only genome doubling, but also the merger of previously divergent evolutionary lineages. Overall, our results suggest that the paleopolyploidy record inferred from current methods is an incomplete account of WGD and reveals deep hybridization preserved in eukaryotic genomes.

---

Polyploidy is a major evolutionary force across eukaryotes and bacteria, with whole-genome duplication (WGD) events documented in fungi, animals, and especially land plants, where ancient polyploidy events have punctuated the phylogenetic history of nearly all major lineages (1–4). In land plants, paleopolyploidy is now understood as a recurrent feature of genome evolution rather than an exceptional condition, with signatures of ancient genome duplication detected across angiosperms, gymnosperms, ferns, and other major clades (5–7). The evolutionary significance of WGD lies partly in the immediate doubling of genetic material, which can relax selective constraints on duplicated genes and provide raw material for functional divergence over time, although most duplicates are ultimately lost (8–10). Ancient WGD has often been treated as both a major source of evolutionary novelty and a central feature of plant genome evolution. Yet the most common evidence for ancient WGD, a peak in the distribution of the number of synonymous substitutions between duplicate genes (9), does not directly record genome doubling, but rather divergence between gene copies that do not recombine.

Detection of ancestral polyploidy events has converged on a simple, elegant method based on patterns in the distribution of synonymous divergence among duplicated genes (9). Specifically, these inferences focus on homologous gene families within a single individual. Once gene families are inferred, pairs of homologous genes are aligned, and substitutions are classified as synonymous or nonsynonymous relative to the number of available synonymous and nonsynonymous sites. A codon-based model is then used to infer synonymous divergence, commonly measured as synonymous substitutions per synonymous site (*K_s_*) (11). Because synonymous substitutions are often treated as approximately neutral, *K_s_* is expected to increase roughly linearly with time after duplicated copies begin evolving independently (9). Therefore, detectable bursts of gene duplication are followed by independent evolution across all duplicated sites, with some gene pairs undergoing decay of *K_s_*.

A *K_s_* peak requires many duplicate gene pairs to begin accumulating mutations at approximately the same time; its detection therefore requires not only duplication, but also divergence, which results from the restriction of recombination or gene conversion between duplicate copies. Thus, *K_s_* peaks are not simple timestamps of genome doubling; they are signatures of when sufficiently divergent gene copies occur in the same genome, either through the union of anciently divergent subgenomes or the accumulation of sequence divergence within a genome through time. This framework has often been explored using birth-death models of gene-family evolution, in which WGD is represented as a pulse of duplicate-gene birth followed by duplicate loss at some rate (12). This generalization has been useful for modeling the expected genomic signature of WGD, but it obscures a key distinction: genome doubling creates additional copies, whereas *K_s_*-based detection requires those copies to exhibit sequence divergence. In allopolyploids, which form via the combined processes of genome doubling and hybridization, most of that divergence may originate before polyploid formation, through prior divergence of the hybridizing lineages.

WGD detection via *K_s_* is biased toward allopolyploids and is less likely to detect autopolyploids, because *K_s_* requires divergence among non-recombining copies. Traditionally, autopolyploids have been defined as having genome copies inherited from within a single species, whereas allopolyploids have been defined as having genome copies inherited from multiple distinct species (reviewed in (14)). Although these classical definitions are often used, they represent only single points along a multidimensional continuum, rather than discrete categories (14). For detectability, however, the more important distinction is mode of inheritance. Allopolyploids are expected to undergo disomic inheritance, where preferential pairing of chromosomes from the same subgenome occurs and crossover between homeologous chromosomes is rare or absent (13). In contrast, autopolyploids are expected to experience polysomic inheritance, where all chromosome copies can pair and there is no preferential pairing. Disomic and polysomic inheritance produce different gametic expectations (Fig. 1), and these inheritance patterns create different evolutionary expectations at both the individual and population levels. In this sense, polysomic autopolyploids initially behave as organisms with “duplicate alleles,” whereas disomic allopolyploids behave as organisms with “duplicate genes” (15). Under polysomic inheritance, duplicated chromosomes continue to pair and recombine, limiting divergence among copies. Under disomic inheritance, subgenomes are isolated, and homeologs are detectable as duplicate genes, with further divergence possible after polyploid formation. Rather than a signature of ancient WGD alone, the *K_s_*-based record of ancient WGD may instead reflect biased hybridization and meiotic isolation. However, this hypothesis remains untested, and we explore it here.

**Fig. 1.**
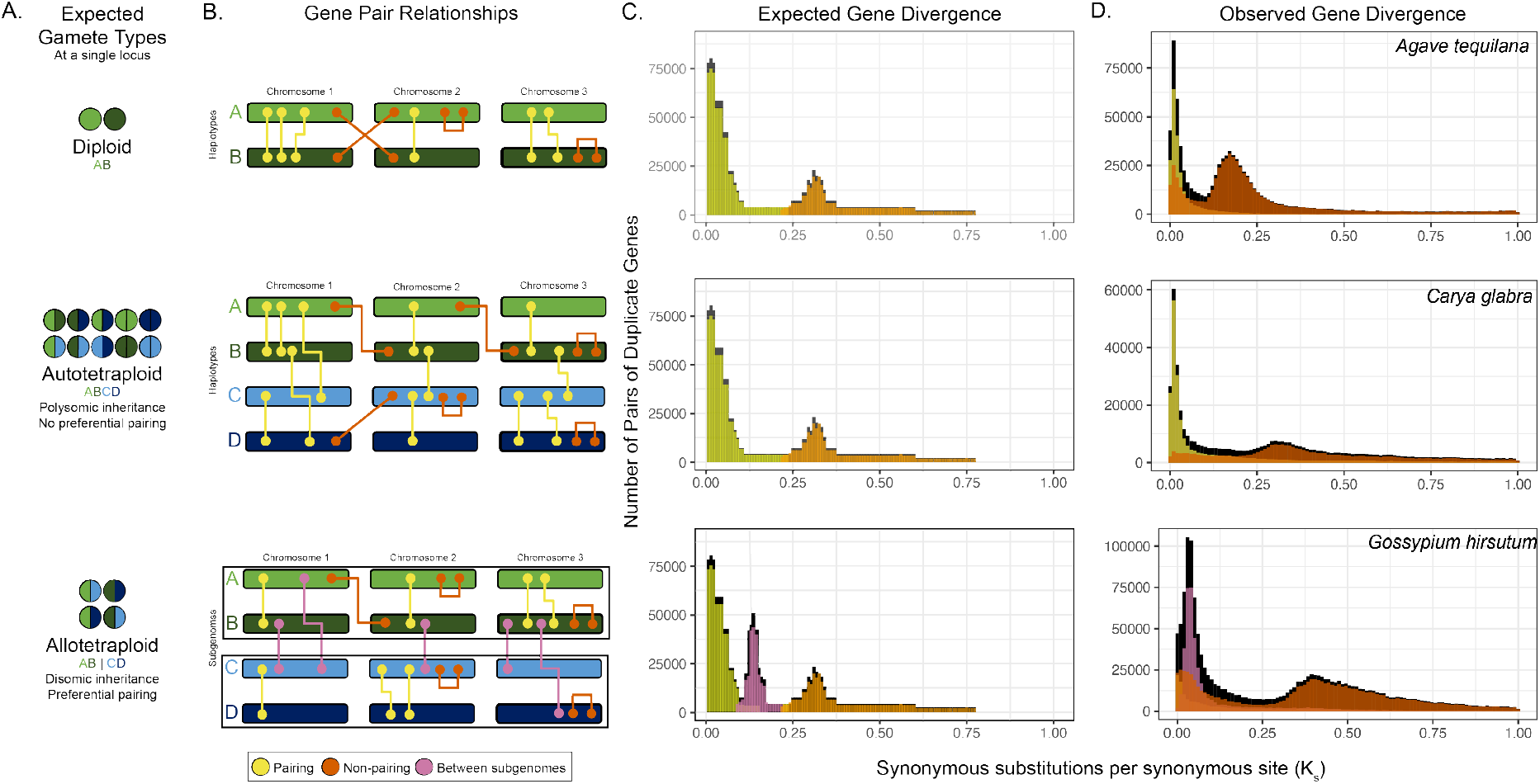
Synonymous substitutions per synonymous site are only accumulated in regions that do not pair during meiosis. A. Diploids, autotetraploids, and allotetraploids have different gametic types expected at a single locus without recombination. In a diploid (AB) gametic types are A or B. For a tetraploid (ABCD), under polysomic inheritance we expect the gametic types AA, AB, AC, AD, BB, BC, BD, CC, CD, and DD. Under disomic inheritance, a tetraploid (ABCD) where AB and CD are homeologs (13), we would expect that gametic types include AC, AD, BC, and BD. B. We can infer *K_s_* values based on the location of a pair of duplicate genes and mode of inheritance. C. We expect gene pairs that occur in pairing regions to be in the exponential and lack synonymous substitutions (yellow). In contrast, gene pairs that do not recombined are able to accumulate substitutions (brown) or accumulate mutations prior to hybridization (pink). D. These predictions are consistent in diploid *Agava tequilana* cultivar Weber’s Blue, autotetraploid *Carya glabra*, and allotetraploid *Gossypium hirsutum* genotype U1 (see SI Appendix, Table S1).

## Results and Discussion

### *K_s_*-based methods detect the absence of recombination between gene pairs

By leveraging haplotype-phased chromosome-scale assemblies of polyploid species (SI Appendix, Table S1 and S2), we directly compared duplicate-gene *K_s_* values with inferred chromosome pairing and inheritance mode (see details in SI Appendix; Fig. 1; SI Appendix, Fig. S1 and S2). Across these assemblies, elevated *K_s_* is concentrated among gene pairs located in chromosomal regions that do not regularly pair or recombine during meiosis (*i*.*e*., inferred as “non-pairing”), whereas gene pairs located on pairing chromosomes or haplotypes show little divergence (*i*.*e*., inferred as “pairing”). This pattern is expected because recombination and gene conversion limit the accumulation of sequence differences between copies. In diploids, detectable synonymous substitutions accumulate between duplicated loci that are not alternative alleles at the same meiotic pairing region, such as duplicates on different chromosomes or duplicated regions within the same haplotype. In an autotetraploid with polysomic inheritance, pairing can occur among all haplotypes (*i*.*e*., A can pair with B, C, or D); therefore, duplicate-gene divergence is restricted to regions isolated from polysomic pairing and recombination. In allopolyploids, by contrast, disomic inheritance separates divergent subgenomes, allowing homeologous gene pairs to retain inherited divergence and accumulate additional substitutions over time. Thus, the empirical pattern across phased assemblies supports the central prediction of our framework. Gene-divergence methods detect chromosome copies isolated from recombination, not genome doubling per se.

For example, in autotetraploid *Carya glabra* (pignut hickory), a peak in the distribution of *K_s_* is restricted to regions inferred as non-pairing and is associated with putative hybridization with *Carya illinoinensis*, rather than with the inferred origin of *C. glabra* 1–3 MYA (Fig 1) (16). In contrast, when disomic inheritance is established in a polyploid, a hybridization-associated signal is more readily retained in duplicate pairs that occur on different subgenomes. In allotetraploid *Gossypium hirsutum* (cotton), gene pairs from different subgenomes have already accumulated synonymous substitutions despite the lineage’s relatively recent origin, approximately 1–2 MYA, similar in timing to the formation of *C. glabra* (Fig 1). Elevated *K_s_* between subgenomes (*i*.*e*., values greater than 0) can therefore reflect divergence that accumulated in the parental lineages before hybridization. Across many haplotype-phased chromosome-scale assemblies, duplicate pairs in non-pairing regions (*i*.*e*., *K_s_* greater than 0) show elevated divergence, whereas pairing regions remain concentrated near low *K_s_* values (SI Appendix, Fig. S1 and S2).

We further explore the fate of autopolyploids and allopolyploids, as well as a sample of their intermediates. We also examine how post-polyploidization processes affect detection of ancestral WGD. We find that, based on current theory and empirical understanding, detection of ancient WGD events is favored when duplicated chromosome copies are already divergent at polyploid formation or subsequently become isolated from recombination. Lastly, we explore the implications of widespread hybridization, rather than WGD alone, across the tree of life.

### Post-polyploidization processess affect detection of ancient WGD

Following WGD, allopolyploids and autopolyploids may both experience multivalent crossover, where more than two chromosomes pair and recombine during meiosis; however, this is not always observed in either allopolyploids or autopolyploids. Multivalent recombination is often considered to be maladaptive because it can increase nondisjunction, produce nonviable or unbalanced gametes, and reduce fertility (but see (17)). Bivalent crossover is increased in canonical allopolyploids through strict preferential pairing, whereas canonical autopolyploids stabilize bivalent pairing through modified crossover dynamics (see below). Although the processes following WGD that lead to stabilized meiosis have collectively been described as “diploidization” per Stebbins (1947) (see SI Appendix) (18), his definition only indicates a return to diploid-like meiosis, or disomic inheritance, in allopolyploids. In contrast, stabilizing meiosis and predominantly bivalent crossover in autotetraploids does not necessarily lead to a shift in inheritance; polysomic inheritance can be present in established autopolyploids (19), and meiosis can be considered stable without preferential pairing (see below). However, transitions from polysomic to disomic inheritance may occur and are explored here.

Allopolyploidization occurs through immediate genome doubling associated with the merging of divergent progenitor genomes. Chromosomal divergence between the parental species can result in immediate preferential pairing. Alternatively, at formation, homeologous pairing and recombination may occur with high prevalence (20). When exchange is reciprocal, homeologous chromosomes may appear as mosaics of the subgenomes. In contrast, when homeologous exchange occurs with replacement, the genome is locally homogenized toward one subgenome (21). Homeologous exchange can lead to shifts to polysomic inheritance (22), as observed in hybrid triploid *Musa acuminata* cultivars where preferential pairing is absent (SI Appendix, Fig. S4; see details in SI Appendix). Prevention of homeologous exchange, or diploidization (per Stebbins (1947); see SI Appendix), can be achieved through strict preferential pairing between homologous chromosomes within each subgenome and limited recombination between homeologous chromosomes. For example, strict preferential pairing may be aided by structural divergence between homeologs; however, control of preferential pairing may be gene-based (reviewed in (15)). Diploidization in allopolyploids is also believed to include gene loss/fractionation, genome downsizing, and chromosomal rearrangements (18). These latter genomic changes may shape the long-term evolutionary consequences of WGD, but they may be distinct from, or reinforce, the initial establishment of independently evolving homeologous copies. The timing of diploidization can be asynchronous across the genome (23, 24), thus altering the signature of WGD in gene-based inferences. Once diploidized, chromosome pairs belonging to each newly formed subgenome can be modeled as distinct, because recombination between them would not occur. Thus, at a population level, allopolyploids can be modeled like two diploid genomes stacked lengthwise.

Autopolyploids have been described as an epigenetic macromutation (25), as they are often accompanied by shifts in phenotype and expression likely due to epigenetic changes and associated shifts in chromatin accessibility (26). For autopolyploids classified as “cytologically diploidized”, meiosis is considered stable, defined by the lack of multivalent crossover (27, 28); however, preferential pairing remains absent, and chromosome copies are expected to pair randomly. In established autotetraploids, bivalent pairing at metaphase I can be established through modified crossover dynamics, rather than through preferential pairing (reviewed in (15, 19); SI Appendix). For autopolyploids, suppression of multivalent crossover may result from shifts in crossover positioning, crossover interference, or a reduction in crossover frequency (15, 27, 29). For example, compared to the diploid progenitor, shortening of chromosomes and/or the synaptonemal complex (the protein structure that links homologous chromosomes during meiosis) ensures a single pair of chromosomes recombine during meiosis despite the lack of preference for which chromosome copies (or haplotypes) pair (15, 29). Additionally, genes that dictate crossover modification may be introduced through subsequent introgression (30). Polysomic inheritance has been observed in polyploids produced through hybridization (31) and in lineages inferred to have hybrid origins (SI Appendix, Table S2). In such cases, homeologous exchange followed by polysomic inheritance could progressively homogenize chromosome copies derived from different progenitors, depending on the recombination rate. If this process removes divergence among chromosome copies, the WGD associated with lineage formation would become difficult to detect using *K_s_*; however, a subsequent shift to disomic inheritance could generate a detectable divergence if chromosome isolation occurs across much of the genome (see below). Compared with disomic inheritance, polysomic inheritance is associated with increased allelic diversity, a higher mutation rate, reduced impact of genetic drift, and relaxed purifying selection relative to a diploid progenitor. Overall, at a population level, autopolyploids must be modeled with all genome copies considered, with the genome copies stacked on top of each other.

The shift from polysomic to disomic inheritance could occur if the duplicated genome is completely purged (*i*.*e*., a complete set of chromosomes is lost) or in instances where preferential pairing evolves; however, only the evolution of preferential pairing would preserve duplicated copies while allowing them to diverge and become detectable in gene-based inferences of ancient WGD. Complete loss of a duplicated genome is likely rare, although it has been observed in snowflake yeast (*Saccharomyces cerevisiae*) (32). Though there is support for the shift to bivalent pairing in polysomic inheritance, the evolution of preferential pairing for polysomic autopolyploids lacks mathematical theory and empirical underpinnings. In theory, genomic rearrangement and/or mutations may allow preferential pairing to evolve. For example, chromosome fusion could promote a shift from polysomic inheritance to disomic inheritance (8). Chromosome fusion has been observed in an autopolyploid snow carp species; both multivalent and bivalent pairing, as well as low genomic divergence (*i*.*e*., 0.00 *< K*_*s*_ *<* 0.05) were identified for these fused chromosomes, interpreted as a shift to disomic inheritance (33). Theoretically, mutations could impact the efficiency of chromosome pairing based on sequence divergence (34, 35) or could occur in specific regions of the genome (35), enabling disomic inheritance. However, when observed in autopolyploids, preferential pairing has been limited to a single chromosome, part of a chromosome, and/or a specific genotype, and is thought to be gene-based rather than driven by broad sequence divergence (36). Preferential pairing would be associated with modified gamete frequencies, based on the location of the locus, but it is unknown whether these signatures could be detected on a macroevolutionary timescale (37). For such a transition to generate a detectable peak in the *K_s_* distribution, the shift from polysomic to disomic inheritance would likely need to occur across a substantial portion of the genome within a relatively narrow evolutionary interval; if shifts are asynchronous, they may be indistinguishable from synonymous divergence due to other processes (see details in SI Appendix) and would not be detectable as peaks in *K_s_* distributions. The extent to which selection would favor a shift to stable disomic inheritance in established autopolyploids with polysomic inheritance - and the conditions under which this might occur - remain unclear (reviewed in (19)).

Despite inheritance type shaping the expectations for chromosome evolution after polyploidization, the presence or absence of preferential pairing does not fully determine structural or average gene divergence between chromosome pairs (Fig. 2; SI Appendix, Fig. S3 and S4). Generally, chromosomal and gene divergence are lower under polysomic inheritance than under disomic inheritance; however, this pattern is not consistent across all species and is also influenced by other factors such as divergence time (Fig. 2; SI Appendix, Fig. S5). In both allopolyploids and autopolyploids, mechanisms that stabilize meiosis can increase recombination between homologous chromosomes relative to their diploid progenitor(s) (38, 39); therefore, substantial gene divergence between regularly pairing homologous chromosomes is not expected under either inheritance type, consistent with the low divergence observed among pairing chromosome copies (Fig. 2). Overall, the outcome of post-polyploidization processes is multidimensional rather than a continuum between two discrete bins.

**Fig. 2.**
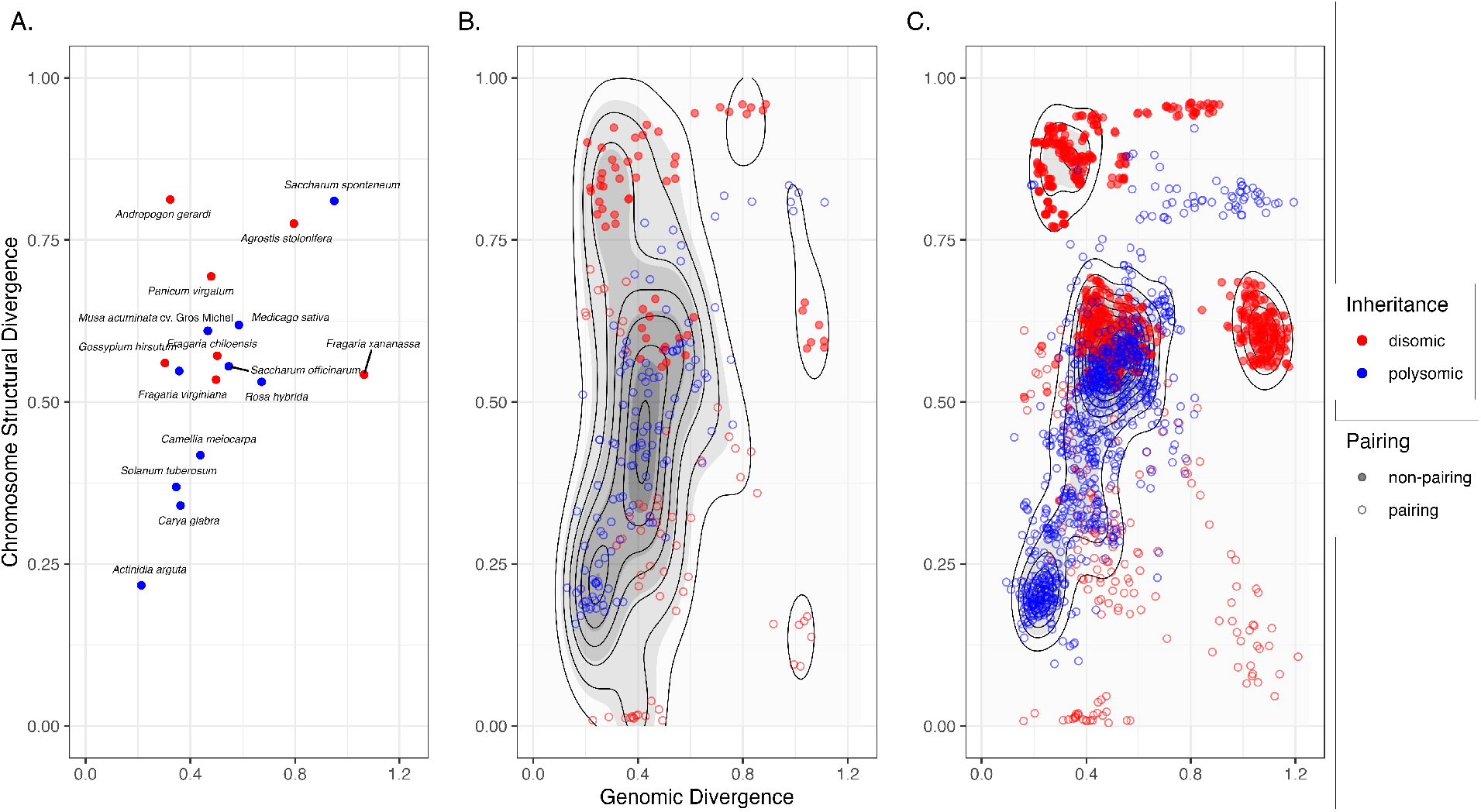
Mode of inheritance does not dictate discrete chromosomal structural divergence or genomic divergence among polyploids. We find chromosomal structural divergence and genomic divergence overlaps among mode of inheritance when A. averaged for each species, B. averaged across haplotypes that have the same pairing expectation for each chromosome, and C. averaged for each haplotype comparison per chromosome. All estimates occurred per chromosome and between haplotypes. Chromosomal structural divergence among haplotypes for each chromosome was inferred based on haplotype assembly graphs. Genomic similarity, or average *K_s_*, was calculated for gene comparisons that occurred only on the same chromosome and on different haplotypes for all assemblies. Generally, chromosomal structural divergence is less in autopolyploids than in allopolyploids; however, chromosomal structural divergence and genomic divergence is generally higher for regions that do not pair during meiosis under disomic inheritance compared to regions that pair under disomic or polysomic inheritance.

### Gene-divergence methods preferentially detect allopolyploids

Considering the microevolutionary consequences and expectations following WGD under disomic vs. polysomic inheritance, we expect gene-divergence methods to preferentially detect polyploids in which duplicated copies have evolved independently over long-term evolutionary timescales (*i*.*e*., disomic inheritance). Paleopolyploidy events are most readily detected when recombination is limited between subgenomes and when divergence either predates polyploid formation through divergence of the parental lineages or accumulates substantially between isolated subgenomes.

Despite the likely lack of detection of genes inherited polysomically in evolutionary time, ancient autopolyploidy events have been suggested in numerous lineages. Some ancient WGD events have been assumed to be of autopolyploid origin because hybridization was considered rare in certain lineages (8). In some instances, such as the teleost-specific WGD, debate remains. Analyses have identified genomic patterns consistent with allopolyploidy (40), as well as patterns that do not fit canonical expectations under either allopolyploidy or autopolyploidy (24). In other cases, for example yeasts, low sequence divergence was attributed to an autopolyploid origin (41), with subsequent analyses supporting an allopolyploid origin (42). Furthermore, genomic sequence divergence is not binary between autopolyploids and allopolyploids (*i*.*e*., genomic divergence overlaps; Fig. 2), so sequence divergence alone should not be used to classify polyploid origin.

Understandings of autopolyploids and allopolyploids have been largely based on disconnected theories; theoretical models applied to allopolyploids are not applicable to autopolyploids and vice versa. For allopolyploids, theoretical frameworks employ birth-death models to examine gene retention over macroevolutionary timescales (*e*.*g*., (12)). For autopolyploids, models focus on population-level dynamics over microevolutionary time periods. Bridging these theoretical frameworks and extending them to incorporate genomic dynamics will be vital for identifying features that could distinguish ancient autopolyploid events. Furthermore, it is important to identify genomic expectations under both types of inheritance, as many ancestral WGD inferences are now based on whole-genome assemblies and subgenome estimation. Models that allow transitions between polysomic and disomic inheritance could provide a framework for predicting when WGD becomes detectable across evolutionary timescales.

Overall, with our current inference and theoretical frameworks, we are likely underestimating the frequency of ancient polyploids because gene-divergence methods are poorly suited at detecting WGD events that retain polysomic inheritance, and thus may systematically miss ancient autopolyploids. The evolutionary importance of polyploidy, as well as the contribution of WGD to modern genome content and structure, may therefore be underestimated.

### The importance of hybridization

If ancient WGDs detected by *K_s_* methods are disproportionately allopolyploid in origin, then the history recovered from peaks in a *K_s_* distribution is not simply a history of genome duplication. It is also a history of hybridization. Genome doubling can occur without producing immediate detectable sequence divergence among duplicated copies, particularly when chromosome copies continue to pair and recombine under polysomic inheritance. Hybridization, by contrast, can unite chromosome sets that have already diverged in separate lineages. Those divergent subgenomes can produce immediately detectable *K_s_* signal, or become detectable over time by remaining isolated by preferential pairing. Thus, detectability depends not only on genome doubling, but also on reticulation, inherited subgenome divergence, and the maintenance and/or evolution of meiotic isolation. Given the current limited evidence for transitions from polysomic to disomic inheritance (see above), the paleopolyploid record may therefore be biased toward events that merged previously divergent genomes while systematically underrepresenting genome duplications that remained polysomic, leading to an overall underestimation of ancient polyploidy.

This reframing also changes the evolutionary question posed by the ancient WGD record. Historically, we have asked how many times genomes doubled. For detectable ancient events, we may also need to ask how many times divergent lineages merged. WGD adds copies, but allele or gene copy number alone does not create the sequence divergence used to identify most ancient polyploid events. Hybridization brings together genomes that have already been evolving independently, creating novel combinations of alleles, regulatory states, chromosome structures, and gene interactions before genome doubling has produced any new divergence of its own. In many allopolyploids, the evolutionary novelty attributed to WGD may therefore derive in part from the merger of divergent genomes, with genome doubling stabilizing, preserving, or amplifying the consequences of that merger (reviewed in (43)).

If detectable ancient WGDs are mostly allopolyploid in origin, several broader implications follow. First, angiosperm history may contain more deep reticulation than is currently represented in strictly bifurcating phylogenies (44, 45). Some regions of the phylogeny associated with ancient WGD may not mark only the divergence of descendant lineages, but also the fusion of previously separated lineages whose genomes were brought together by hybridization (reviewed in (44)). Second, hybridization may have contributed to lineage persistence during periods of environmental upheaval by combining divergent genetic backgrounds, increasing standing variation, or facilitating the movement of adaptive variants across lineages (see (46– 48)). These possibilities do not imply that all major nodes are reticulations, or that the tree model is uninformative. Rather, they suggest that some key nodes in plant evolutionary history may be better understood as points where divergence and reticulation co-occur, with ancient WGD signals preserving evidence of both.

Modern phylogenomics often treats hybridization, WGD, and incomplete lineage sorting (ILS) as separate processes. But over tens of millions of years, genomes fractionate, chromosomes rearrange, and duplicates are lost, while accumulating substitutions can obscure the signal needed to distinguish hybridization from incomplete lineage sorting. Eventually, one of the most persistent signals of hybridization may be a duplicate-gene divergence peak. Ancient hybridization may be easier to detect in polyploid descendants, because polyploidy can preserve divergent parental gene copies long after other signatures of reticulation have been obscured or lost.

Lineages with abundant inferred ancient WGDs should also show evidence of recurrent hybridization, subgenome structure, or other genomic signatures consistent with the merger of divergent genomes. Conversely, ancient autopolyploid events should be difficult to detect using gene-divergence-based methods unless ancient autopolyploids subsequently evolve mechanisms that isolate chromosome copies and permit near-simultaneous divergence. Ancient autopolyploids may, however, become detectable through approaches that more directly model polysomic inheritance, allele dosage, chromosome pairing, or other population-genomic signatures of genome doubling.

Viewed through the lens of our analyses, the paleopolyploid record is not merely a record of genome duplication. Instead, it may be one of the deepest and most pervasive account of hybridization preserved in eukaryotic genomes.

## Materials and Methods

We briefly describe data inclusion and analysis; detailed materials and methods are in the SI Appendix.

### Genomic data inclusion

We obtained and analyzed publicly available haplotype-resolved, chromosome-scale genome assemblies based on inclusion criteria (see details in the SI Appendix). Overall, we identified and analyzed 17 genome assemblies for taxa ranging from 3*x* to 8*x* and spanning eight orders of angiosperms (SI Appendix, Table S1).

The included genome assemblies had various naming schemes for genes, haplotypes, and chromosomes. Therefore, for each assembly, we manually created genome assembly keys that identified relationships among genes, haplotypes, and chromosomes. For species identified as allopolyploids, genome assembly keys included subgenome identity. For all analyses, we used chromosome identity and haplotype identity as the genome assembly coordinate system. For example, the presence of two genes on the same chromosome does not indicate whether they occur on the same or different haplotypes (*i*.*e*. copies of that chromosome). Similarly, two genes on the same haplotype do not necessarily share the same chromosome. For visualization, see Figure 1.

### Gene divergence inference

For each genome assembly, coding DNA sequences were concatenated from all haplotypes, gene families were then inferred, and synonymous substitutions per synonymous site (*K_s_*) were calculated between every gene pair in a gene family. To create painted *K_s_* distributions, we filtered and partitioned the resulting pairwise comparisons (see details in SI Appendix). Briefly, for each assembly, we used the genome assembly key to classify each pairwise gene comparison, or gene pair, into an expected pairing category based on chromosome, haplotype, and, when available, subgenome identity (see Fig. 1). These categories describe whether the chromosomal regions containing each gene pair are expected to align and potentially recombine during meiosis (see SI Appendix for more information).

### Similarity and inheritance inference

We constructed haplotype assembly graphs for each chromosome, including all haplotypes regardless of subgenome identity. We measured chromosomal structural similarity among haplotypes for each genome assembly (see details in the SI Appendix). Lastly, we assessed each chromosome’s inheritance pattern; we used properties of each genome assembly to classify inheritance as disomic or polysomic (see SI Appendix for details).

## Data, Materials, and Software Availability

No data were generated for this study. All genome assemblies are publicly available.

## ACKNOWLEDGMENTS

The authors thank Alyssa Phillips (Atlanta Botanical Garden) and Shengchen Shan (University of Florida) for assistance identifying assembled genomes. We thank Jacob S. Berv (University of Michigan) for feedback on the manuscript draft. MLG was funded through a National Science Foundation Postdoctoral Research Fellowship in Biology (DBI-2410238). P.S.S. and D.E.S. were funded in part by National Science Foundation grant DBI-2320251. S.A.S. and K.F. were funded in part by National Science Foundation grants IntBio-2217116, DEB-NERC-1939226, and DBI-1930030.

