## Supplemental Appendix 1 for "All detectable ancient whole-genome duplications involve hybridization"

#### This PDF file includes:

Supporting text
Figs. S1 to S5
Tables S1 to S3
SI References

### Supporting Information Text

### Methods

**Genomic data inclusion.** Haplotype-resolved, chromosome-scale assemblies with FASTA files available for all assembled haplotypes, Coding DNA Sequence (CDS) identified with an associated FASTA and/or annotation file (*i.e.*, General Feature Format (GFF) and/or Browser Extensible Data (BED)), and known ploidal level were identified and manually downloaded from a diversity of publicly available databases (Table S1). All genome assemblies were downloaded by June 15, 2026. For most published genome assemblies, phased haplotypes or annotations were not available; therefore, the number of genome assemblies that could be included was limited. In total, we identified and analyzed 17 genome assemblies for taxa that ranged from $3x$ to $8x$ and spanned eight orders of angiosperms (Table S1). We also included a genome assembly for a species (*Agave tequilana* cv. Weber's Blue) now considered a diploid but reported to have undergone a putative ancestral whole-genome duplication (WGD) (1). The included genome assemblies had various naming schemes for genes, haplotypes, and chromosomes. Therefore, for each assembly, we manually created genome assembly keys that identified relationships among genes, haplotypes, and chromosomes.

Note, chromosome and haplotype identities are used as the coordinate system of a genome assembly. For example, the presence of two genes on the same chromosome does not indicate whether they occur on the same or different haplotypes (*i.e.* copies of that chromosome). Similarly, the presence of two genes on the same haplotype does not indicate that these genes are found on the same chromosome. For visualization, see Figure 1.

**Gene divergence inference.**

***Inferring $K_s$.*** We inferred synonymous substitutions per synonymous sites ($K_s$) for gene pairs for all included species. For each species, CDS regions were sampled from each haplotype assembly, and the resulting sequences were merged into a single FASTA file. With this FASTA, we used wgd v.2.0.28 (2) to infer gene families and paralogous gene pairs (`wgd dmd`). This approach defines gene families based on clustering (2). Pairwise synonymous substitutions per synonymous site ($K_s$; `wgd ksd --pairwise`) were then calculated for every gene pair. Although pairwise estimates are susceptible to oversaturation (3, 4), they were needed to classify gene pairs into pairing categories (described below).

Genome assembly keys were then used to identify whether pairwise gene comparisons occurred between genes on the same or on different haplotypes, as well as between genes on the same or on different chromosomes. For species identified as allopolyploids, subgenomes were first identified based on the primary literature associated with the assembly (Table S1; Table S2) and assessed using inheritance pattern analyses (described below). For these species, genome assembly keys identified if gene pairs occurred on the same or different subgenomes. For all subsequent analyses, comparisons were excluded when either gene lacked haplotype identity or chromosome identity.

***Painting $K_s$.*** To create painted $K_s$ plots, we first filtered pairwise $K_s$ estimates to remove comparisons with $K_s < 0$, $K_s > 1$, or missing $K_s$ values. We then assigned each gene pair to an expected pairing category based on chromosome, haplotype, and, when available, subgenome identity. These categories describe whether the chromosomal regions containing each gene pair are expected to align and potentially recombine during meiosis. They do not represent direct measurements of recombination or meiotic pairing. The pairing categories are visually represented in Figure 1. As mentioned above, chromosome and haplotype identities are used as the coordinate system of a genome assembly (see Figure 1).

For diploids and autopolyploids, gene pairs were categorized as "pairing" if they occurred on the same chromosome and on different haplotypes. These regions are considered homologous chromosomes and are expected to align during meiosis and may recombine. Gene pairs were categorized as "non-pairing" when located in regions not expected to pair during meiosis, including pairs on different chromosomes regardless of haplotype identity and pairs located on the same chromosome within the same haplotype. For allopolyploids, which all had assigned subgenomes, gene pairs were categorized as "pairing" when located on the same chromosome, different haplotypes, and the same subgenome (*i.e.*, homologous chromosomes). Gene pairs categorized as "non-pairing" included pairs on different chromosomes regardless of haplotype or subgenome identity, as well as pairs located on the same chromosome, same haplotype, and same subgenome. We also defined a third category, "non-pairing between subgenomes," for gene pairs located on the same chromosome, different haplotype, but different subgenomes, where preferential pairing is not expected (*i.e.*, homeologous chromosomes).

We expected "pairing" gene pairs to have low $K_s$ values because ongoing recombination or gene conversion should limit the accumulation of sequence divergence between copies. We did not estimate local recombination rates, which are not available for all included assemblies and are known to vary across plant genomes (5), likely as a function of structural variation, chromatin state, and epigenetic regulation (6–8). Moreover, suppression of recombination can occur through many mechanisms (8). Therefore, some gene pairs classified as "pairing" may still accumulate synonymous substitutions. However, such cases are expected to broaden the low-$K_s$ distribution of pairing regions rather than produce a synchronous duplicate-gene peak associated with WGD.

We expected "non-pairing" gene pairs to exhibit higher $K_s$ values (*i.e.*, values greater than 0) because the chromosomal regions containing them do not regularly recombine. For allopolyploids, divergence between progenitor species creates historical sequence divergence between subgenomes before polyploid formation. Gene flow between progenitor lineages or homeologous exchange after WGD could reduce this divergence, whereas strict preferential pairing should preserve or increase divergence between subgenomes through time.

We examined the resulting pairwise $K_s$ values for all included genome assemblies (Fig. S1). To verify that pairwise $K_s$ did not lead to misinterpretation due to oversaturation, we calculated mean $K_s$ for each gene family based on pair type

(Fig. S2). Both pairwise and gene-family analyses supported the expectations, where "pairing" gene pairs were found in the exponential and concentrated near low $K_s$ values, whereas signatures of older divergence were concentrated in "non-pairing" and "non-pairing-between-subgenomes" categories.

#### Similarity and inheritance inference.

***Haplotype graph construction.*** For each genome assembly, we constructed a haplotype assembly graph (9) for each chromosome consisting of haplotypes from a single individual. To construct the haplotype graph, we manually constructed FASTA files for each chromosome, containing complete chromosomes from every haplotype. The resulting FASTA files were then indexed with samtools v.1.20 [`faidx`; (10)]. The haplotype graphs were then created using pggb v.0.7.4 with haplotype number specified, minimum average nucleotide identity for segments set to 90%, and scaffold length set to 5000 (9).

***Similarity inference.*** Chromosomal structural similarity was measured among haplotypes for each genome assembly. These similarity metrics were obtained based on the resulting odgi file for each chromosome with odgi v. 0.9.4 [`odgi similarity`; (11)]. First, the Jaccard index was estimated between haplotypes, where the shared base pairs between the two haplotypes, or intersection, is divided by the sum of each haplotype's total length in base pairs minus the intersection (*i.e.*, $jaccard = \frac{intersection}{(hapA+hapB-intersection)}$). Using the estimated Jaccard distance, estimated identity was obtained equal to $\frac{(2*jaccard)}{(1+jaccard)}$. We then converted this estimate to a measure of chromosomal structural divergence among haplotypes, by subtracting the estimated identity from one. To match this estimated identity for our pairwise $K_s$ comparisons, we filtered the $K_s$ comparisons to include only pairs that occur on the same chromosome and different haplotypes (see above). To remove $K_s$ estimates that are likely due to saturation, the comparisons were filtered to remove any value where $K_s$ was greater than 5 or less than 0. We also removed comparisons where $K_s$ was not calculated. Average $K_s$ was then calculated for the complete genome assembly, for each chromosome divided into pairing types, and for each haplotype comparison at each chromosome.

***Inheritance inference.*** To infer inheritance of each chromosome, we assessed patterns consistent with disomic or polysomic inheritance using properties of each genome assembly. Here, we used the haplotype graphs defined above where all haplotypes for each chromosome were aligned. We then extracted biallelic sites from these alignments; in the resulting genotype, one allele is from each haplotype.

Prior to inferring inheritance, for each species, we investigated if mode of inheritance (*i.e.*, disomic or polysomic) had been identified based on both parental and progeny sequencing. Many of the species for which genome assemblies are included here have confirmed inheritance patterns, although inheritance is often inferred for a different cultivar or genotype (see Table S2). Here, we leveraged confirmed inheritance to verify our inheritance inference framework, focusing on *Solanum tuberosum* and *Gossypium hirsutum*. Polysomic *Solanum tuberosum* is an exemplar model, as inheritance was confirmed for the same cultivar as the reference genome (12). In contrast, disomic *Gossypium hirsutum* is a well-known allopolyploid evolutionary model (13). Additionally, the framework used here to infer inheritance is based on fundamental concepts (14, 15), which have previously been applied when inferring and modeling inheritance (16–18). As introduced in the main text, disomic and polysomic inheritance will result in different gamete types. At a biallelic locus, for a tetraploid (AABB), under polysomic inheritance we expect the gametic types AA, AB, and BB (*i.e.*, no preferential pairing; see Table S3). Under disomic inheritance, with preferential pairing between homeologous chromosomes within each subgenome, a tetraploid (AABB) with AA and BB representing homeologs will produce only AB gametes. Alleles within each subgenome will segregate independently of other subgenomes, and homeologous loci will exhibit fixed heterozygosity (17, 19).

To assess whether inheritance is disomic or polysomic, biallelic single nucleotide polymorphisms (SNPs) were examined for each haplotype alignment graph. For simplicity, we focus on biallelic loci as it is very difficult to correct and/or account for alignment and variant calling errors with more than two alleles per site. The graph file (GFA) was first deconstructed with vg v.1.74.1 (20) with sites included from all snarls and the reference set as the primary haplotype (`vg deconstruct -a -P HapA`). The resulting Variant Call Format (VCF) file was filtered with bcftools v.1.23.1 (10) to only biallelic sites (`bcftools view --types snps -m 2 -M 2`). The filtered VCF was then converted to a tab-delimited file with grep (`grep -v '^##'`). From each VCF, we isolated the reference allele, alternative allele, and variant call for each haplotype (where reference allele = 0, alternative allele = 1). For every genome assembly, we then examined (1) the frequency of each heterozygote type, and (2) haplotype allelic identity (Fig. S3 and Fig. S4), described below.

To investigate the frequency of heterozygote types, we first calculated the frequency of the alternative allele (*i.e.*, B) for each SNP. For example, in a tetraploid, we expect sites at $AAAB$ (B frequency = 0.25), $AABB$ (B frequency = 0.50), and $ABBB$ (B frequency = 0.75). Along each chromosome, the frequency of each heterozygote type was then calculated. Under disomic inheritance, it is expected that the majority of SNPs will occur at $AABB$ (*i.e.*, $AABB > AAAB + ABBB$) due to fixed heterozygosity between subgenomes. Because we only include biallelic SNPs, we expect the majority of these loci to differ between subgenomes rather than within a subgenome, which would be expected due to segregating homologous loci. Under polysomic inheritance, we expect gametes of AA, AB, and BB at biallelic loci. Unlike disomic inheritance, we do not expect random assortment of haplotypes to shift genotype frequencies towards a specific genotype; based on Hardy-Weinberg Equilibrium, the probability of each heterozygote is equal to $P(AAAB) = 4p^3q$, $P(AABB) = 6p^2q^2$, and $P(ABBB) = 4pq^3$ (15) (see Table S3). At heterozygous sites, where the alternative allele frequency is greater than zero and less than one and has a uniform distribution, we expect relatively equal proportions of $AAAB$, $AABB$, and $ABBB$ (17). For our tetraploid genome assemblies, each chromosome was classified as disomic when $AABB > (AAAB + ABBB)$ and polysomic when $AABB < (AAAB + ABBB)$ (Fig. S3). These patterns are consistent for both polysomic *Solanum tuberosum* and disomic *Gossypium hirsutum* (Fig S3). Although graph-based alignments mitigate reference bias (*i.e.*, bias towards the reference nucleotide compared to the alternative nucleotide) by allowing paths between sites, we only retained sites where the reference and alternative contained a single nucleotide; therefore, reference bias is expected, and the relative proportion of $AAAB > AABB > ABBB$ is possible. Additionally, for higher ploidal levels, due to increased types of heterozygotes, it is difficult to identify modes of inheritance based on frequency of heterozygote types alone. Additionally, population-level processes, like selection, can shift these genotype frequencies away from the expectation.

To provide an additional assessment of inferred inheritance patterns and preferential pairing, we also examined haplotype allelic identity. For each chromosome, we recorded the proportion of SNPs where allelic identity was equal between two haplotypes, and this process was repeated between every pair of haplotypes. Without preferential pairing, we expect equal proportions between haplotype pairs, as observed in polysomic *Solanum tuberosum* (Fig S4). When preferential pairing occurs, we expect the haplotype allelic identity proportion to be higher between preferentially paired haplotypes than between non-preferential pairs. In a tetraploid, under strict disomic inheritance with divergent subgenomes, the proportion of haplotype allelic identity between preferential haplotypes (*i.e.*, "pairing") should be larger than the proportion of haplotype allelic identity between non-preferential haplotypes (*i.e.*, "non-pairing"); this representation allows us to account for potential segregating homologous loci. This pattern is consistent for disomic *Gossypium hirsutum* (Fig S4). Notably, this metric is a measure of two haplotypes having the same identity along a chromosome, rather than two haplotypes sharing the same identity with only each other along a chromosome; proportions calculated for each pair do not add to one, because this measure is not considering allelic identity with additional haplotypes. Therefore, when subgenomes are similar to each other, the proportion of haplotype allelic identity will be higher. All comparisons were examined per chromosome, as labeling often does not indicate relatedness among chromosomes. For example, units labeled haplotype A chromosome 1 and haplotype A chromosome 2 may actually belong to different haplotypes.

Measures of inheritance, here based on the frequency of heterozygote types and haplotype allelic identity, should be interpreted carefully and skeptically. At the per-site level, deviations can be due to genome misassembly, missing data in the genome assembly, and/or haplotype graph inference error. Here, depth of each inference is limited to the number of haplotypes, limiting our ability to detect error in the assembly and/or biological processes such as double reduction and homeologous exchange. Ideally, raw genomic sequencing data or population-level sequencing data should be mapped back to each reference to confirm inheritance patterns. For example, rather than relying on only the assembled haplotype-phased genome, *Solanum tuberosum* inheritance was confirmed with cytology and population sequencing of both parental and progeny genotypes, which identified a large proportion of multivalent formation and presence of double reduction (12). Although the lack of preferential pairing and polysomic inheritance is confirmed for *Solanum tuberosum* based on our inference (Fig. S3 and Fig. S4), the large variance in our inference of haplotype allelic identity should not be interpreted as indicative of a biological process without additional analysis and investigations. Similarly, the presence of homeologous exchange should not be inferred without additional analysis (21).

Notably, the classifications of four genome assemblies did not match the inferred mode of inheritance (Table S2): *Musa acuminata* cv. Cavendish, *Musa acuminata* cv. Gros Michel, *Rosa hybrida*, and *Camellia meiocarpa*. *Musa acuminata* cv. Cavendish and *Musa acuminata* cv. Gros Michel are classified as allotriploids due to their origin via hybridization (22); however, homeologous exchange with replacement is hypothesized to have led to shifts in haplotype composition and led to the lack of preferential pairing observed here [see Fig. S4; (23, 24)]. For tetraploids *Rosa hybrida* and *Camellia meiocarpa*, an allotetraploid origin was mentioned, but neither assembly included the designation of subgenomes. Subgenome identification is inferred through comparisons with potential progenitors (*i.e.*, extant diploid species) and clustering-based approaches, often focused on long terminal repeat retrotransposons, an abundant transposable element in plants. For *Rosa hybrida*, haplotypes were found to be mosaics of multiple progenitors likely due to extensive homeologous exchange (25). For other *R. hybrida* cultivars, preferential pairing is found only for certain genotypes (26). For *Camellia meiocarpa*, haplotypes were found to be distinct from *C. olerifera*, but conclusions about origin were not made (27). Based on phylogenetic data, *C. meiocarpa* has been suggested to have originated from two closely related diploid species (28).

### Supplementary Text

***Diploidization.*** "Diploidization" was initially introduced in terms of haploid mycelium becoming diploid [(29); see pg. 8 in (30)]. In relation to polyploids, Stebbins (1947) introduced the term "diploidization" by contrasting young vs. established allopolyploids. In young allopolyploids, the presence of "heterogenetic associations" (30), or the pairing of chromosomes from divergent subgenomes, is likely to result in sterility (31–33). In contrast, in established allopolyploids, the absence of pairing of chromosomes from divergent subgenomes, or strict disomic inheritance, would indicate progression to the diploid (*i.e.*, disomic) state. Notably, Darlington (1937) defined "heterogenetic associations" as the segregation of dissimilar chromosomes, which may occur with bivalent or multivalent formation [see pg. 206 in (30)]; therefore, this does not refer simply to the formation of bivalents or multivalents (*i.e.*, a cytogenetic pattern), but an inheritance or segregation pattern. Stebbins (1947) described a "diploidized" allopolyploid *Nicotiana* (34) with normal meiosis, classified by bivalent formation, preferential pairing, and lack of homology between subgenomes, likely due to the elimination of duplicate genes. For polyploids, bivalent formation is now referred to as "cytological diploidization", while lack of homology due to the elimination of duplicate genes is referred to as "genic diploidization" (35).

Based on Stebbins's (1947) definition of "diploidization", this term likely was only meant to apply to allopolyploids with disomic inheritance, supported by the belief at the time that autopolyploids were evolutionary dead-ends (31). Despite the intended use of "diploidization" for only allopolyploids, the available data led to the application of this term based on the observation of chromosome pairing for both disomic and polysomic inheritance. Specifically, diploidization was assessed by the presence or absence of multivalents during metaphase I and by the fertility of the resulting offspring (36–38), rather than the presence or absence of preferential pairing of chromosomes. As reviewed by Bomblies (2023), autopolyploids that undergo polysomic inheritance, and therefore have chromosomes that randomly pair during meiosis, can also form bivalents at metaphase I [*e.g.*, (39–44)], despite potential multivalent formation during prophase I. Therefore "cytogenetic diploidization" does not indicate a return to a diploid-like state or the absence of multivalent formation; rather, the term implies the absence of multivalent crossover at metaphase I, and the resulting genomic stability provided by predominantly bivalent crossover. Furthermore, "cytogenetic diploidization" does not imply a single path to genomic stability or a specific mode of inheritance (45). Although bivalent pairing is often referred to as diploid-like, it does not indicate disomic inheritance. The transition from polysomic segregation to disomic segregation [*i.e.*, the allopolyploidization of an autopolyploid; (46)] would require bivalent pairing with non-random partners.

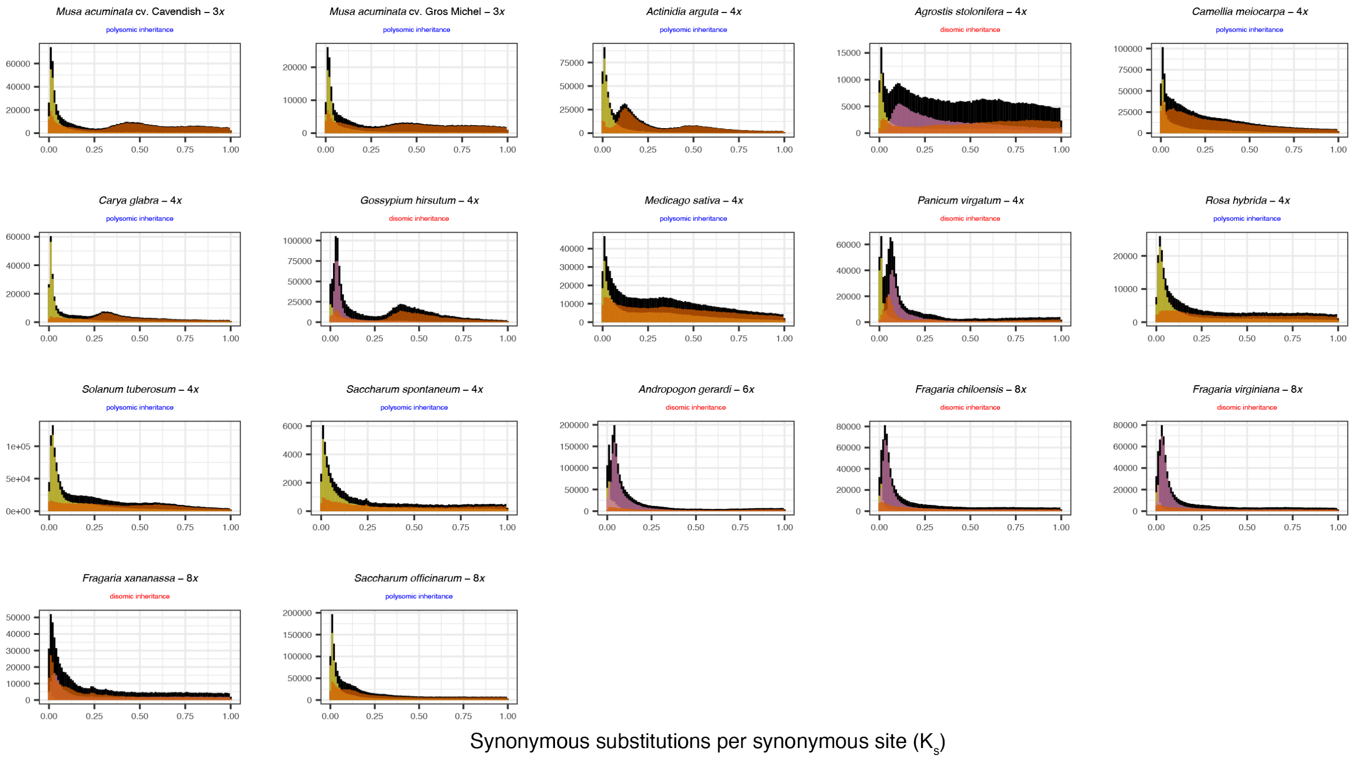


**Fig. S1.** Across many polyploid species, synonymous substitutions per synonymous site are only accumulated in regions that do not pair during meiosis. Here, all genes pairs (black) are classified into three classes: (1) pairing (yellow): gene pairs that occur on different haplotypes and same chromosome, therefore expected to align during metaphase (2) non-pairing (orange): gene pairs that occur in regions that are not expected to align during metaphase, (3) non-pairing between subgenomes (pink): gene pairs that occur on haplotypes that do not preferentially pair during meiosis, classified only in allopolyploid taxa. For genotypes and cultivars designations see Table S1.

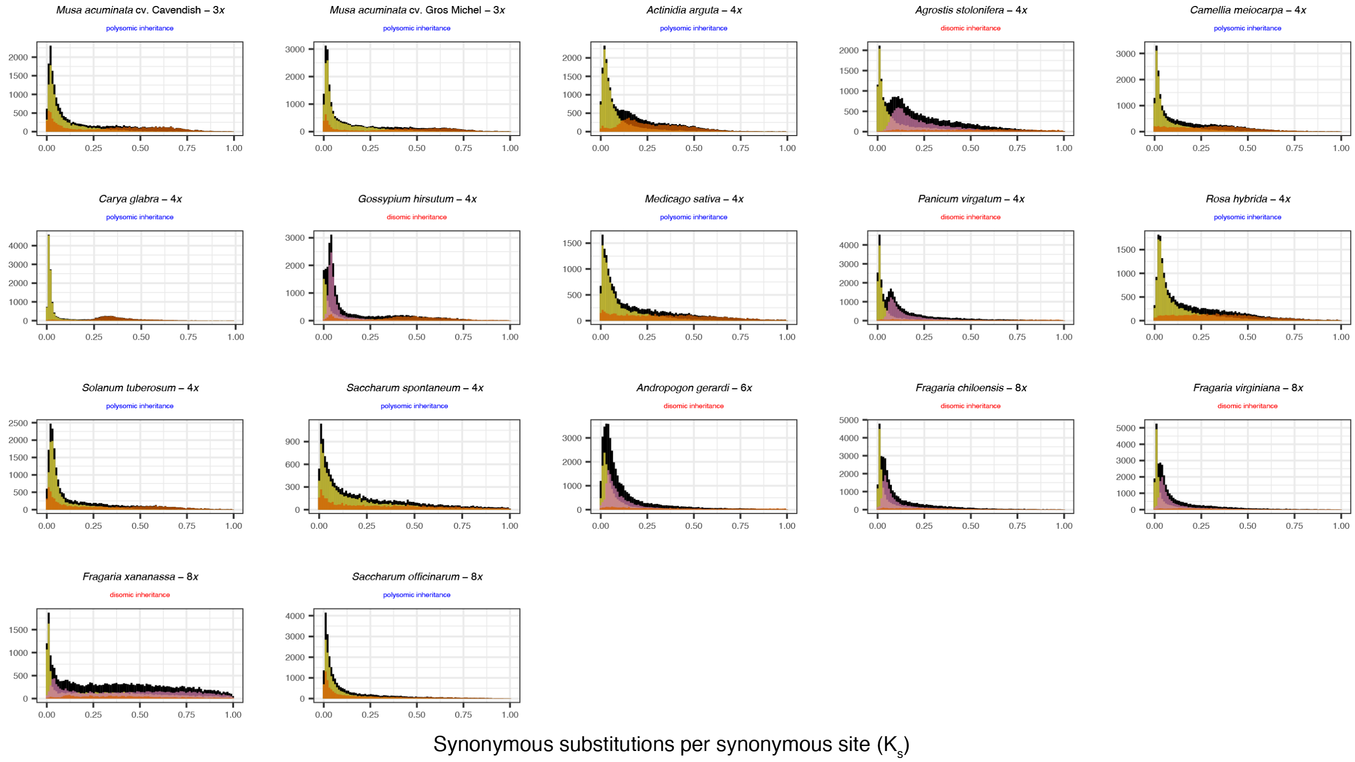


**Fig. S2.** Average $K_s$ per gene family for each pairing type is consistent with pairwise $K_s$ estimates. Synonymous substitutions per synonymous site are only accumulated in regions that do not pair during meiosis. Here, all genes pairs (black) are classified into three classes: (1) pairing (yellow): gene pairs that occur on different haplotypes and the same chromosome, therefore expected to align during metaphase (2) non-pairing (orange): gene pairs that occur in regions that are not expected to align during metaphase, (3) non-pairing between subgenomes (pink): gene pairs that occur on haplotypes that do not preferentially pair during meiosis, classified only in allopolyploid taxa. For genotypes and cultivars designations see Table S1.

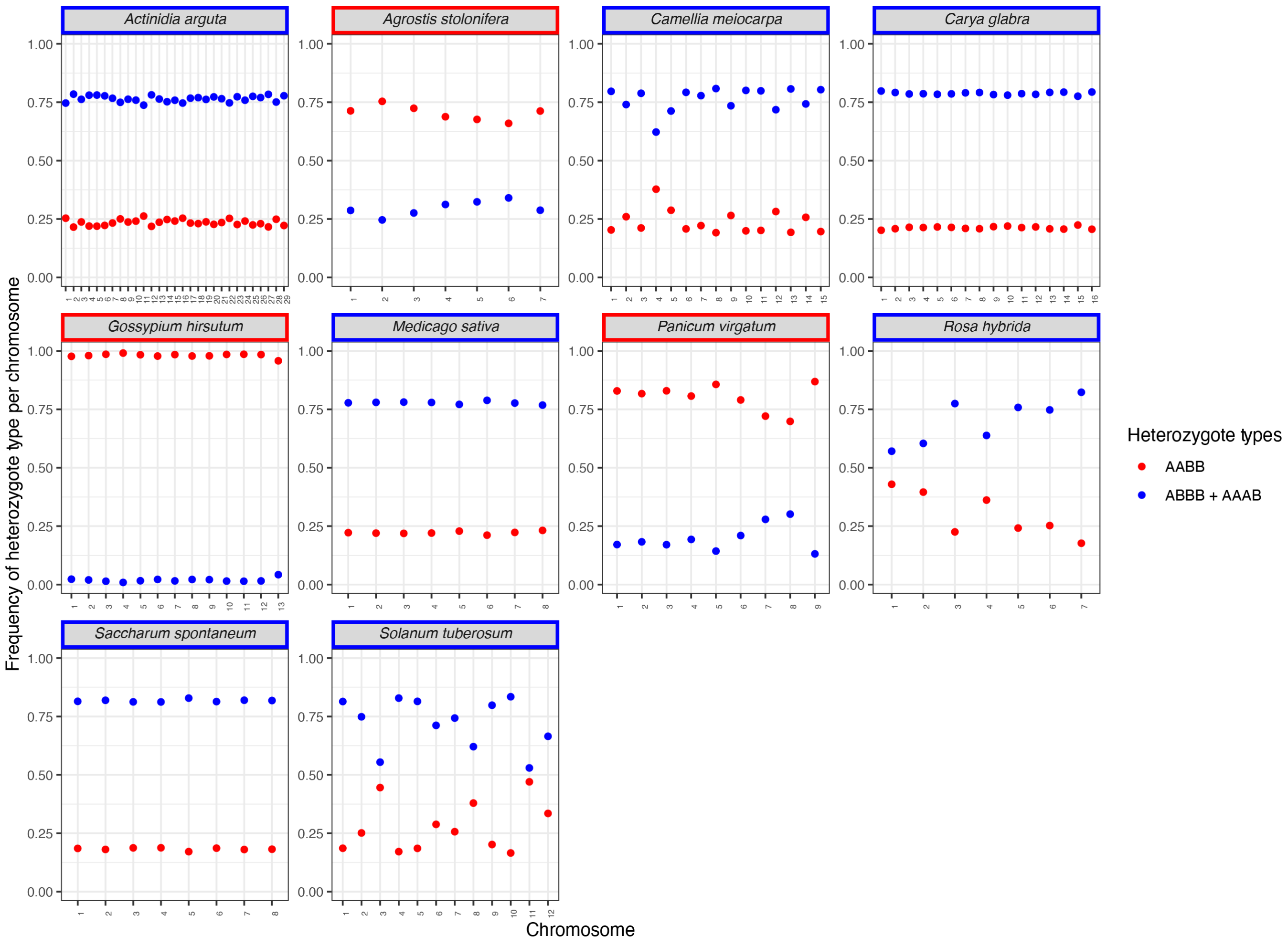


**Fig. S3.** Site-based heterozygosity indicates inheritance type for tetraploid assemblies. To determine if assembly represents disomic inheritance (red) or polysomic inheritance (blue), we investigated biallelic variants extracted from haplotype graphs. With disomic inheritance, one expects fixed heterozygosity where all copies from the same subgenomes have the same allele (ex. $AABB$, see *Gossypium hirsutum* var U1). With polysomic inheritance (blue), greater proportions of $AAAB + ABBB$ (blue points) are expected compared to $AABB$ (red points), while the opposite is expected under disomic inheritance. See Figure S4 for ploidal levels greater than 4*x*. For genotypes and cultivars designations see Table S1.

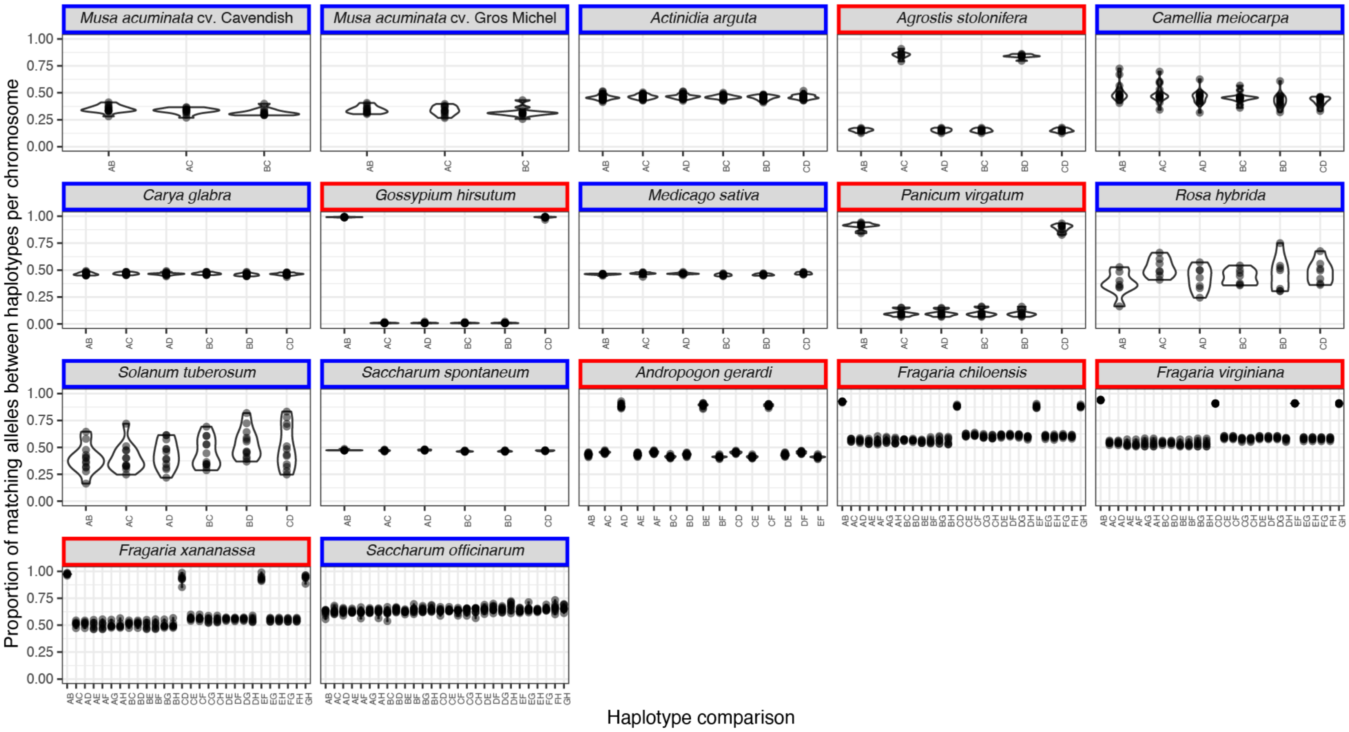


**Fig. S4.** For all ploidal levels, preferential pairing is obvious when the proportion of sites that have matching alleles between two haplotypes is counted across biallelic SNPs for each chromosome. Under disomic inheritance (red) when preferential pairing is ongoing, the proportion of sites that match in allelic identity between the preferential haplotypes is higher than between non-preferential haplotypes. Therefore, we expect two clusters for each chromosome under disomic inheritance and for these clusters to be relatively consistent across chromosomes; the difference between these two clusters can be minimal (ex. *Fragaria* spp.) or discrete (ex. *Gossypium hirsutum*). Comparatively, under polysomic inheritance (blue), haplotypes do not preferentially pair and the proportion of shared alleles should be relatively similar between haplotype pairs. For genotypes and cultivars designations see Table S1.

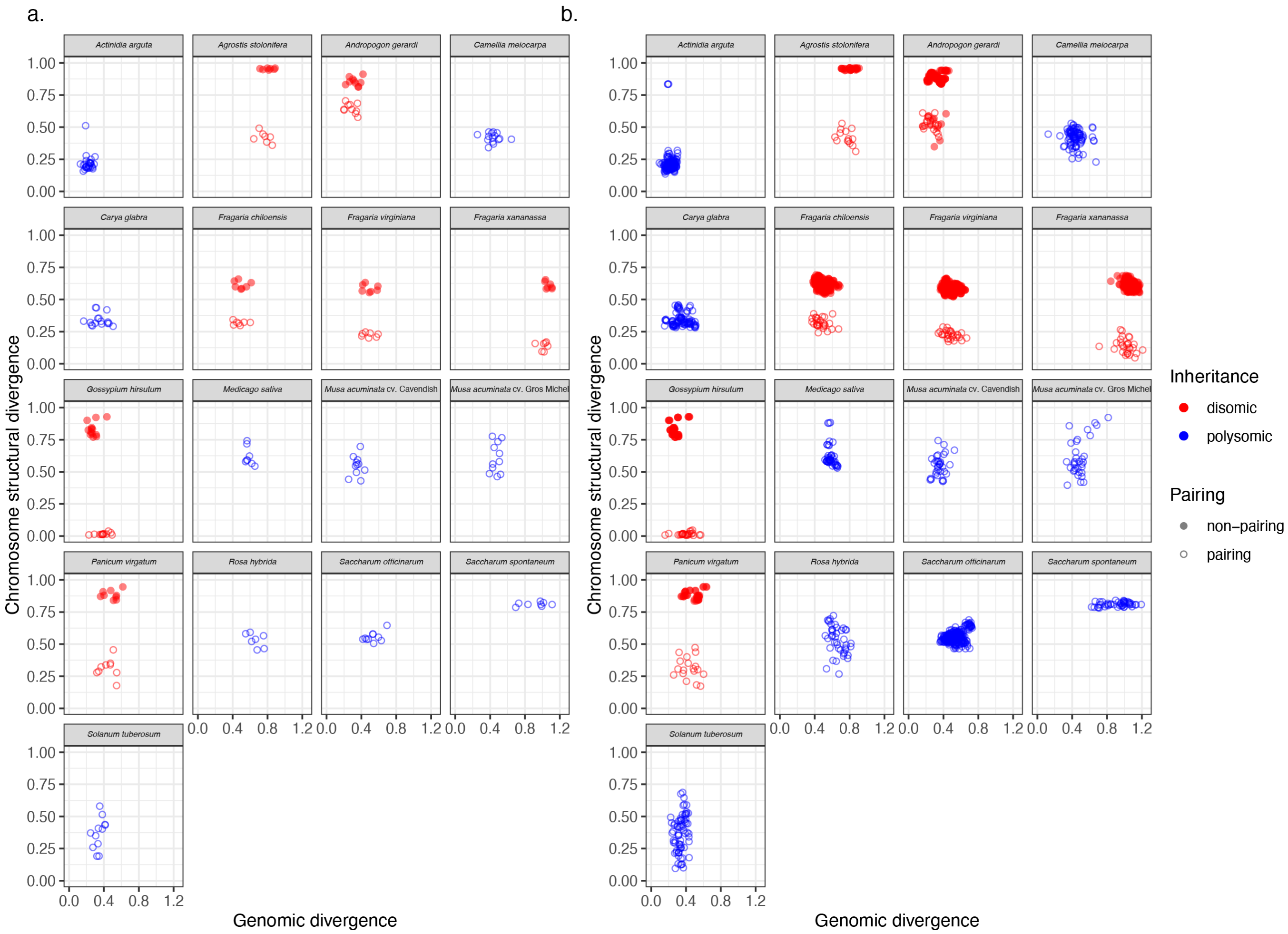


**Fig. S5.** Disomic inheritance occurs with distinct chromosomal divergence among pairing and non-pairing chromosomes. Following Figure. 2, chromosomal structural divergence and genomic divergence was A. averaged across haplotypes that have the same pairing expectation for each chromosome, and B. averaged for each haplotype comparison per chromosome. Chromosomal structural divergence among haplotypes for each chromosome was inferred based on haplotype assembly graphs. Genomic divergence, or average $K_s$, was calculated for gene comparisons that occurred only on the same chromosome and on different haplotypes for all assemblies. Under polysomic inheritance, clusters do not form among haplotypes and chromosomal structural divergence is similar among all haplotype pairs. For genotypes and cultivars designations see Table S1.

**Table S1. Genome assemblies included in any analysis. Scientific names match the name assigned to the genome assembly and include taxonomic authorities, as well as genotype and cultivar designations, as relevant. Data repositories from which data were downloaded include the United States Department of Energy's Joint Genome Institute (DOE-JGI), National Center for Biotechnology Information (NCBI), United States Department of Agriculture Database (USDA; gdatacommons.nal.usda.gov), China's National Genomics Data Center (NGDC), Dryad (https://datadryad.org/), FigShare (http://figshare.com/), and taxon-specific repositories.**

| Scientific Name | Family | Order | Ploidal Level | Data Repository | Citation |
|---|---|---|---|---|---|
| *Agave tequilana* F.A.C.Weber cv. Weber's Blue | Asparagaceae | Asparagales | 2*x* | DOE-JGI | (47) |
| *Musa acuminata* Colla cv. Cavendish | Musaceae | Zingiberales | 3*x* | FigShare | (22) |
| *Musa acuminata* Colla cv. Gros Michel | Musaceae | Zingiberales | 3*x* | FigShare | (22) |
| *Actinidia arguta* (Siebold and Zucc.) Planch. ex Miq. cv. M1 | Actinidiaceae | Ericales | 4*x* | NGDC | (48) |
| *Agrostis stolonifera* L. cv. Declaration | Poaceae | Poales | 4*x* | NCBI and USDA | (49) |
| *Camellia meiocarpa* Hu | Theaceae | Ericales | 4*x* | FigShare | (27) |
| *Carya glabra* (Mill.) Sweet | Juglandaceae | Fagales | 4*x* | NCBI | (50) |
| *Gossypium hirsutum* L. genotype U1 | Malvaceae | Malvales | 4*x* | DOE-JGI | (51, 52) |
| *Medicago sativa* L. cv. XinJiangDaYe | Fabaceae | Fabales | 4*x* | FigShare | (53) |
| *Panicum virgatum* L. genotype WBC | Poaceae | Poales | 4*x* | DOE-JGI | (54, 55) |
| *Rosa hybrida* (Ser.) Posp. cv. Samantha | Rosaceae | Rosales | 4*x* | FigShare | (25) |
| *Solanum tuberosum* L. cv. C88 | Solanaceae | Solanales | 4*x* | solomic.agis.org.cn | (12) |
| *Saccharum spontaneum* L. cv. AP85-441 | Poaceae | Poales | 4*x* | sugarcane-genome-hub.southgreen.fr | (56) |
| *Andropogon gerardi* Vitman | Poaceae | Poales | 6*x* | JGI-DOE | (57) |
| *Fragaria chiloensis* (L.) Mill. | Rosaceae | Rosales | 8*x* | rosaceae.org | (58) |
| *Fragaria virginiana* Duch. | Rosaceae | Rosales | 8*x* | rosaceae.org | (58) |
| *Fragaria xananassa* Duch. cv. Florida Brilliance | Rosaceae | Rosales | 8*x* | rosaceae.org | (59) |
| *Saccharum officinarum* L. cv. Purple | Poaceae | Poales | 8*x* | sugarcane-genome-hub.southgreen.fr | (60) |

**Table S2. Inheritance and classification of each included polyploid. Classification and hybrid origin are reported based on primary literature; see references in Table S1. Progeny Inference indicates inferred inheritance based on a progeny array for this species; note, cultivar or genotype may differ between progeny inference and assembled genome. Mode of Inheritance indicates inheritance inferred based on genome assemblies for all polyploid reference genomes (See Fig. S3 and Fig. S4; see (? )). Asterisks indicate instances where the Mode of Inheritance based on genome assembly, as implemented here, did not match the classification assigned in the primary literature; see supplemental methods for more information on these species. For genotypes and cultivars designations, see Table S1.**

| Scientific Name | Ploidal Level | Classification | Hybrid Origin | Progeny Inference | Mode of Inheritance |
|---|---|---|---|---|---|
| *Musa acuminata* cv. Cavendish | 3*x* | Allotriploid | Yes | - | Polysomic* |
| *Musa acuminata* cv. Gros Michel | 3*x* | Allotriploid | Yes | - | Polysomic* |
| *Actinidia arguta* | 4*x* | Autotetraploid | Unlikely | Mostly Polysomic (61) | Polysomic |
| *Agrostis stolonifera* | 4*x* | Allotetraploid | Yes | Disomic (62) | Disomic |
| *Camellia meiocarpa* | 4*x* | Allotetraploid | Suspected | - | Polysomic* |
| *Carya glabra* | 4*x* | Autotetraploid | Unlikely | - | Polysomic |
| *Gossypium hirsutum* | 4*x* | Allotetraploid | Yes | Disomic (63) | Disomic |
| *Medicago sativa* | 4*x* | Autotetraploid | Unlikely | Polysomic (64) | Polysomic |
| *Panicum virgatum* | 4*x* | Allotetraploid | Yes | Disomic (65, 66) | Disomic |
| *Rosa hybrida* | 4*x* | Allotetraploid | Yes | Mostly Polysomic (26, 67) | Polysomic* |
| *Solanum tuberosum* | 4*x* | Autotetraploid | Yes | Polysomic (12) | Polysomic |
| *Saccharum spontaneum* | 4*x* | Autotetraploid | Unlikely | Mostly Polysomic (68, 69) | Polysomic |
| *Andropogon gerardi* | 6*x* | Allohexaploid | Yes | - | Disomic |
| *Fragaria chiloensis* | 8*x* | Allooctoploid | Yes | Disomic (70) | Disomic |
| *Fragaria virginiana* | 8*x* | Allooctoploid | Yes | Disomic (70) | Disomic |
| *Fragaria xananassa* | 8*x* | Allooctoploid | Yes | Disomic (70) | Disomic |
| *Saccharum officinarum* | 8*x* | Autooctoploid | Unlikely | Polysomic(68) | Polysomic |

**Table S3. Hardy-Weinberg equilibrium for autotetraploids and allotetraploids. Here, p is the frequency of the reference allele and q is the frequency of the alternative allele. For an autotetraploid, these probabilities are the result of $(p+q)^4$. For an allotetraploid with disomic inheritance, we must consider the frequency of the reference allele and alternative allele at two loci; therefore we consider the frequency of alleles at locus A ($p_A$ and $q_A$) and at locus B ($p_B$ and $q_B$). The probabilities for an allotetraploid are the result of $(p_A+q_A)^2 * (p_B+q_B)^2$.**

| Genotype | Autotetraploid Probability | Allotetraploid Probability |
|---|---|---|
| AAAA | $p^4$ | $p_A^2 p_B^2$ |
| AAAB | $4p^3q$ | $2p_A^2 p_B q_B + 2p_A q_A p_B^2$ |
| AABB | $6p^2q^2$ | $p_A^2 q_B^2 + q_A^2 p_B^2 + 4p_A q_A p_B q_B$ |
| ABBB | $4pq^3$ | $2p_A q_A q_B^2 + 2q_A^2 p_B q_B$ |
| BBBB | $q^4$ | $q_A^2 q_B^2$ |
